# Policy lessons from spatiotemporal enrollment patterns of Payment for Ecosystem Service Programs in Argentina

**DOI:** 10.1101/421933

**Authors:** Mauricio M. Núñez-Regueiro, Lyn C. Branch, Josh Hiller, Cristina Núñez Godoy, Sharmin Siddiqui, José Volante, José R. Soto

## Abstract

Over the last 50 years, payment for ecosystem services schemes (PES) have been lauded as a market-based solution to curtail deforestation and restore degraded ecosystems. However, PES programs often fail to conserve sites under strong long-term deforestation pressures and allocate financial resources without having a sizeable impact on long-term land use change. Underperformance, in part, is likely due to adverse selection as landowners with land at the lowest threat from conversion or loss may be most likely to enroll or enrollment may be for short time-periods. Improving program performance to overcome adverse selection requires understanding attributes of landowners and their land across large scales to identify spatial and temporal enrollment patterns that drive adverse selection. In this paper, we examine these patterns in Argentina’s PES program in the endangered Chaco forest ecoregion, which was established in 2007 under the National Forest Law. Our study area covers 252,319 km^2^. Among our most important findings is that large parcels of enrolled land and land owned by absentee landowners show greater evidence of spatiotemporal adverse selection than smaller plots of land and land owned by local actors. Furthermore, lands managed for conservation and restoration are more likely to be associated with adverse selection than lands that provide financial returns such as harvest of non-timber forest products, silviculture, and silvopasture. However, prior to recommending that PES programs focus on land uses with higher potential earnings, a greater understanding is needed of the degree to which these land uses meet ecological and biodiversity goals of PES programs. Because of this, we posit that a PES incorporating a market-based compensation strategy that varies with commodity prices, along with approaches that provide incentives for conservation and restoration land uses and enrollment of local landowners, could promote long-term conservation of endangered lands.

## 1. Introduction

Current deforestation levels threaten biodiversity and provision of ecosystem services such as CO_2_ storage and mitigation of climate change. Incentive-based strategies such as Payments for Ecosystem (or Environmental) Services (PES) programs have been championed as a means to preserve forests by providing financial incentives to forest owners who voluntarily enroll in payment schemes and, in exchange for payments, commit to continued provision of ecosystem services (Wunder et al., 2008). However, effectiveness of PES may be hindered when participants enroll land with low threat of deforestation (i.e., lands that would likely remain forested regardless of PES), thus increasing the budgetary cost of the program while having only a minimal impact on deforestation (adverse selection; Ferraro, 2008). An analogous situation occurs when land under high deforestation threat is enrolled for short time periods. These projects may provide little return for conservation but still increase the cost of the PES program (adverse selection in time, Drechsler et al., 2017; Núñez-Regueiro et al., in review). Temporal and spatial enrollment patterns are critical for understanding this issue because they drive adverse selection, one of the greatest challenges for PES (Ferraro, 2008; Sims et al. 2014; Pagiola et al., 2016; Drechsler et al., 2017).

Factors that may contribute to spatial or temporal enrollment patterns and result in adverse selection are diverse, including characteristics of landowners and landholdings, and restrictions on land uses under the PES program. For example, landowners who depend primarily on agricultural production for their livelihood might be less likely to enroll highly productive lands, and thus lands under higher threat of deforestation, than landowners who either have other sources of income or manage land for purposes other than income generation (Alix-Garcia et al., 2015; Robinson et al. 2016; Silva et al., 2016). Similarly, local landowners closely involved with agricultural production may be less willing to enroll lands for long periods of time than absentee landowners, who generally have other sources of income (Miranda et al., 2003; Arriagada et al., 2009; Petrzelka et al., 2013). Landholding size also may influence enrollment. Large landowners may have smaller agricultural production cost, higher profits per unit of land, and thus less incentive to enroll land with high potential for agriculture than small landowners (Cushman, 2006; Arriagada et al., 2012). The land use allowed on a property under PES also could influence spatial and temporal patterns of enrollment, and thus probability of adverse selection (Arriagada et al., 2009; Miteva et al., 2012). More restrictive land uses may reduce profits and limit the ability of landowners to track markets, and thus discourage long term enrollment of highly productive lands.

To date, studies of adverse selection have focused on whether participants enroll lands that are threatened, but the importance of contract length has received little attention (Arriagada et al., 2009; Alix-Garcia et al., 2015; Sims and Alix-Garcia, 2017), possibly because most PES programs have a fixed contract length (e.g., four-five years, Schomers and Matzdorf, 2013; Ezziene-de-Blas et al., 2016; Pagiola et al., 2016; Sims and Alix-Garcia, 2017). However, even with fixed contracts, renewable contracts provide opportunities for variable enrollment times for landholdings. Moreover, studies of PES that include information on attributes of landowners and their land are rare (Miteva et al., 2012; Schomers and Matzdorf, 2013; Ezziene-de-Blas et al., 2016). Thus, understanding of how characteristics of the landholding, landowner attributes, and land use relate to adverse selection remains limited.

In this paper, we identify predictors of spatial and temporal enrollment patterns in Argentina’s PES scheme. We focus on the PES program in the Chaco dry forest in northwestern Argentina, a forest with one of the highest levels of deforestation worldwide fueled by rapid expansion of soybean and livestock production (Grau et al., 2008; Hansen, 2013; Volante et al., 2016; Nolte et al. 2017a; Fehlenerg et al., 2017). Like other PES programs, this one aims to avoid deforestation through voluntary enrollment of threatened land (Schomers and Matzdorf, 2013; Ezziene-de-Blas et al., 2016; Pagiola et al., 2016; Sims and Alix-Garcia, 2017). A key characteristic that distinguishes the Argentine PES program from others is that contract lengths vary among participants (García Collazo et al., 2013). This, in combination with readily available information on agricultural potential and other characteristics of landholdings, economic activities of landowners, and land use, provides a rich platform to identify factors that contribute to both temporal and spatial adverse selection.

We hypothesized that the primary economic activity of landholders would be a strong driver of temporal and spatial enrollment patterns linked to adverse selection. Participants in this PES program range from private landowners with large agricultural operations to real estate agents, indigenous communities, and local governments managing public land (Graziano Ceddia and Zepharovich, 2017; le Polain de Waroux et al., 2017; Marinaro et al., 2017). In addition to primary economic activity of landowners, we examined three other factors that could influence enrollment patterns related to adverse selection: 1) whether landowners were local or absentee, 2) size of enrolled property, and 3) type of land-use allowed at the site under PES, which ranged from partial forest removal for silvopastoral management to strict conservation.

Overall, we found that primary economic activity was not the most parsimonious predictor of adverse selection, though this was related to other factors. Alternatively, the land uses allowed, whether landowners were local or absentee, and size of enrolled parcels were better predictors of observed spatial and temporal adverse selection. Our findings suggest that the Argentine PES program will have stronger chances of enrolling land with high agricultural potential and longer contracts if it targets local landowners who submit small parcels and land-use plans that have the potential to provide additional earnings, such as silviculture, silvopasture, or non-timber forest products. Relatively few large parcels were enrolled, particularly in areas of high deforestation threat, and enrollment was shorter for large parcels, which represents a fundamental challenge for both this program and environmental conservation of the threated Chaco.

## 2. Methods

## 2.1 Study Area

The Chaco forest of Argentina, Bolivia, and Paraguay is the second largest forested ecoregion in South America, after the Amazon, and a key global conservation area because of high levels of biodiversity and endemism (Fehlenberg et al. 2017; Kuemmerle et al., 2017). The Chaco suffers from record-high land-conversion driven by large-scale agriculture (Grau et al., 2008; Gasparri et al. 2013; Kuemmerle et al., 2017). Sixty percent of the Chaco occurs in Argentina, and many rural and indigenous communities rely on the natural resources of this region. Our study focuses on the four provinces in Northwestern Argentina that hold the largest tracts of Chaco forest: Chaco, Formosa, Salta, and Santiago del Estero (252,319 km^2^, Fig. 1). This landscape consists of remnant forest strips, small forest patches, and some larger forest blocks embedded in a matrix of large farms primarily used for soybeans or cattle pasture (Núñez-Regueiro et al., 2015).

**Fig. 1.**
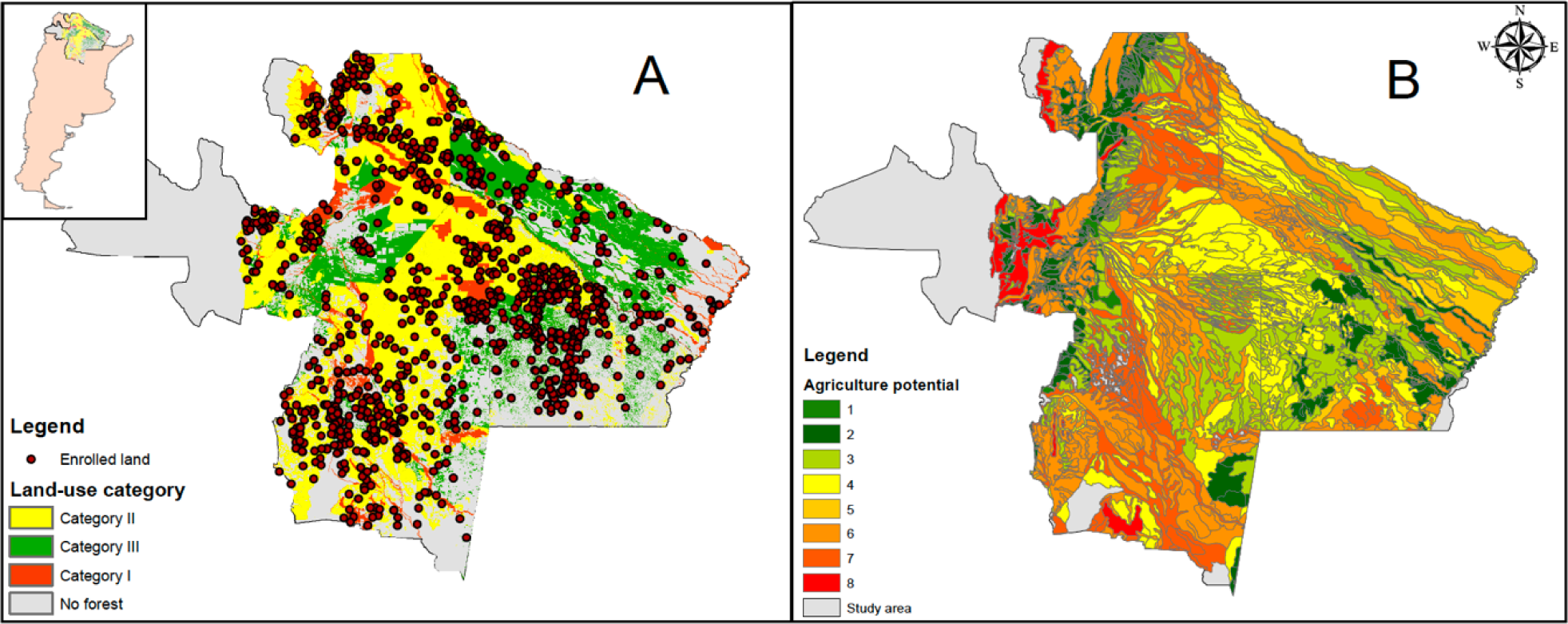
A. Study area comprised of the four provinces that contain most of the Chaco forest in Argentina and corresponding land-use zoning categories for this forest, ranging from high protection to low protection (red, yellow, and green). B. Agricultural suitability index for the study area. Highest suitabilities are presented with darker red colors. Abbreviation codes: L.U. plan type = Land-use type (conservation, restoration, non-timber forest/silviculture, or silviculture/silvopasture) Local/Absentee. = Local or absentee landowner Size = Size of parcel enrolled in PES Primary activity = Primary economic activity (Government/nonprofit, campesinos or indigenous communities, row crop agriculture, cattle ranching, silviculture, real estate or other non-agricultural business). DF = Degrees of freedom logLik = Log-Likelihood AICc = Akaike Information Criterion corrected for finite sample sizes Delta AICc = difference in AICc between best model and each individual model Weight = model weight (Akaike weight)

## 2.2 Program Description

In recognition of the need to curb high rates of deforestation, Argentina’s legislature passed an innovative law (Native Forest Law 26331; henceforth Forest Law) in 2007 that established a minimum annual federal budget for environmental protection, enrichment, restoration, conservation, and sustainable management of native forests and the environmental services they provide (García Collazo et al., 2013, Aguilar et al. 2018). Under the Forest Law, the federal government sets general requirements for environmental protection and funds a national-level PES program that distributes funds to the provinces. Each province defines its own conservation priorities and management objectives, and decides how to allocate the funds (Nolte et al., 2017a). The Forest Law required national-level land-use planning to identify, prioritize, and protect important land for local communities and biodiversity conservation. Each province with native forest was responsible for classifying forest into three categories (Fig. 1): Category I (red zone) designates areas of very high value that cannot be deforested or selectively logged. Lands in this zone can be used for conservation, restoration, collection of non-timber forest products, or eco-tourism. Category II (yellow zone) contains areas of medium or high value that should not be deforested but can be used sustainably. Landowners can choose any land use in category I as well as silviculture and silvopasture. Category III (green zone) represents areas of low value that can be cleared of forest. If landowners with land in this category choose to enroll in the program, they have the same land-use choices as landowners with land in the other two categories. The Forest Law stipulates financial compensation for each participating province based on the amount of land in each land-use category and to individual program participants who voluntarily enroll their land in the payment program (García Collazo et al., 2013). The administrative unit for this program is the cadastral parcel. A landowner can own several parcels but only enroll one or a few parcels, or even part of a single parcel. Participants first enroll each parcel, or portion of a parcel, for one year by submitting a “formulation project” aimed at collecting baseline biodiversity information. Then, participants submit a plan (in Spanish, “certificado de obra”) for each parcel where landowners commit to conduct activities to maintain or enhance ecosystem services, according to the land uses allowed under each zoning category, and define the contract length for the parcel. Payments are conditional on landowners providing this plan.

Arguably, establishing payments in areas with high land-use restrictions provides little additionality (i.e., payments might be redundant given that the Forest Law already protects land in red and yellow categories), and evidence regarding effectiveness of this program as a mechanism to decrease deforestation is the subject of current debate (Nolte et al., 2017a and 2018; Volante and Seghezzo, 2018). Untangling relative contributions of land-use restrictions and payments schemes for reducing deforestation is difficult. However, recent studies show that under the Forest Law, deforestation occurs in restricted areas (i.e., red and yellow zones; Camba Sans et al., 2018) and provincial governments have difficulties enforcing the law (Volante and Seghezzo, 2018). Offering payments in areas with land-use restrictions could serve two purposes:1) to compensate landowners for provincially-mandated land-use restrictions and thus, increase overall acceptance and likelihood of compliance with the Forest Law, and 2) provide enrollees with financial resources to sustainably manage their land (Native Forests Law 26331).

Between 2010 – 2015, the PES program allocated over US$45 million to 1,341 projects in the four Chaco provinces (Núñez-Regueiro et al., in review). This investment resulted in almost 43,000 km^2^ of land enrolled (equivalent to 17% of available land that could potentially be enrolled in all zoning categories). The geographic scope of Argentina’s PES program in the Chaco is among the largest in the world (Ezziene-de-Blas et al. 2016; le Polain de Waroux et al., 2017). See Supporting Information for more details.

## 2.3 Data Sources and Variables

To assess how characteristics of enrollees and the land enrolled relate to spatiotemporal enrollment patterns, we first obtained a list of geo-referenced properties enrolled in the first five years of the PES program in Chaco (n = 1,341; 2010-2015) from the PES database of the Argentine Ministry of Environment. This dataset also provided the length of contractual obligations, parcel size, size of land enrolled (i.e., part or all of the parcel), proposed land-use activity under PES, and individual participant identification number. Using this identification number, we then searched on an open-access governmental database with self-reported information on the primary revenue-generating activity of individuals or organizations (henceforth, primary activity). Although self-reported information can introduce bias into datasets, this problem should be minimum in our study because the Federal Administration of Public Income requires individuals to document their claims and monitors for accuracy.

We classified participants into one of seven categories. The first category included governmental and other non-profit organizations that primarily manage land for conservation (e.g., parks and reserves), rather than for commercial purposes, but also manage other fiscal lands that may be leased to private individuals or companies for a fee. The second category comprised indigenous and campesino (peasant) communities that own a mix of revenue-generating lands (e.g., lands for agriculture and cattle ranching) and forested land used for subsistence-level natural resource extraction. The remaining landowners (individuals or companies) were divided into five categories based in their primary economic activity as follows: row crop agriculture (hereafter agriculture), cattle ranching, silviculture, legal/real estate, and other non-agricultural businesses (https://seti.afip.gob.ar/padron-puc-constancia-internet/ConsultaConstanciaAction.do, https://www.cuitonline.com). This database also provided information on city of residence for each landowner. We defined absentee landowners as participants with a PES project in a province different from the landowners’ province of residence. For our final database, we discarded all projects without information on the primary activity of participants and projects listed as “formulation projects,” as these projects are one-year projects that are meant to lead to longer contracts (final number of projects used = 762).

Then, we defined spatial and temporal adverse selection for program participants based on contract length and agricultural suitability of enrolled properties. Spatial adverse selection can increase as productivity of enrolled land decreases (Ferraro et al., 2011). For analysis, we defined high spatial adverse selection as enrollment of land in sites with less than median regional potential agricultural productivity, based on enrolled and non-enrolled lands (Hi-SAS, Fig. 2). Sites in Chaco forest with low productivity potential for agriculture are less likely to be deforested than high-productivity sites (Grau et al., 2008; Fehlenberg et al., 2017) and thus, may be more likely to remain forested in the absence of program interventions (Ferraro et al., 2011). We obtained agricultural suitability for enrolled areas from a raster grid database of the land-use suitability of the region for soybean and pasture for cattle (Fig. 1; data from the National Institute of Agricultural Technology; see Supporting Information for details).

**Fig. 2.**
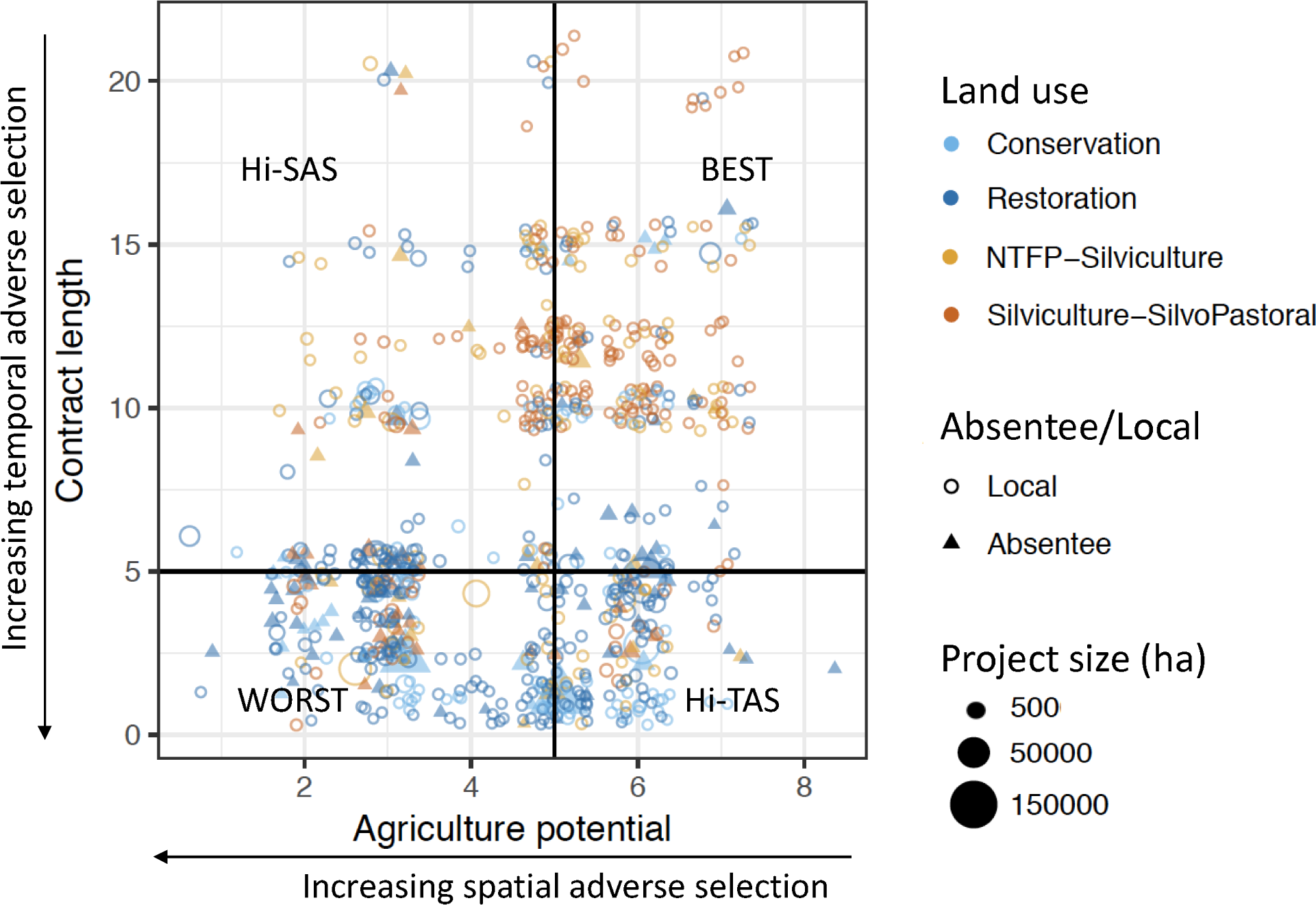
Distribution of enrollment of land with different land-use plan types along axes of increasing temporal and spatial adverse selection. Vertical and horizontal lines represent median agricultural potential and median contract length, respectively. The four quadrats defined by these median values correspond to the four categories of spatiotemporal adverse selection used in our analysis: spatial and temporal self-selection (WORST), primarily high spatial adverse selection (Hi-SAS), primarily temporal adverse selection (Hi-TAS), and no adverse selection (BEST).

Temporal adverse selection is evidenced when threatened land is enrolled for short time periods, thus, reducing funds for long-term contracts (Núñez-Regueiro et al., in review). We defined high temporal adverse selection as enrollment of land in PES for less than the median regional enrollment length (Hi-TAS, Fig. 2). Considering the high deforestation levels in Chaco and the long time-periods required for forest regrowth, short-term contracts likely provide little protection and thus are unsuited to secure long-term provision of environmental services (Grau et al. 2008, Lennox and Armsworth 2011, Drechsler et al. 2017, Fehlenberg et al., 2017; Núñez-Regueiro et al., in review). The Argentine PES program allows participants to re-enroll (García Collazo et al., 2013). However, the degree to which participants will choose this option is unknown. Because the program is fairly new (Forest Law passed in 2007; first participants enrolled in 2010), insufficient data were available to include re-enrollment in our analysis. If substantial re-enrollment occurs, temporal adverse selection could be lower than reported in our study.

Depending on the agricultural production potential where a given PES project is located and its contract length, a participant was categorized as only having high spatial or high temporal adverse selection (Hi-SAS or Hi-TAS, respectively), both high spatial and high temporal adverse selection (WORST), or low spatial and temporal adverse selection (BEST). Avoiding both temporal and spatial adverse selection would require enrolling land under high threat of conversion for long periods of time (Fig. 2).

## 2.4 Data Analysis

To test our hypothesis and understand other factors that may contribute to spatial and temporal enrollment patterns, we built a series of multinomial logistic regression models (MLRM) with spatiotemporal adverse selection as a response variable with the four categories (Hi-SAS, Hi-TAS, BEST, and WORST). To understand the relationship between adverse selection and primary activity of participants, we used a categorical predictor variable: levels one trough five corresponded to the five revenue-generating activities listed above, level six to land management by governmental and non-governmental organizations, and level seven to land management by peasant and indigenous communities. To test competing hypotheses, we also included the following predictor variables: absentee or local landowner (binary variable), size of land enrolled (continuous numerical variable), and type of land-use plan enrolled (categorical variable with four levels: conservation plan, restoration plan, silviculture and collection of non-timber forest products, and silviculture and silvopastoral activities). The government database did not separate silviculture from non-timber forest products and silvopastoral activities or list ecotourism as a land use. The size of the land enrolled corresponds to the size of the parcel or portion of parcel that was enrolled. For convenience we will refer to this as parcel size. The total landholding size of landowners was not recorded in the PES database, and thus is unknown. To reduce multicollinearity in our model set, we tested for associations between variables and only included variables with no statistical evidence of associations among potential predictors in the same model (see Table 1 for list of models and Table S-1 for correlations among variables).

**Table 1.** Multinomial logistic regression model comparison for spatiotemporal adverse selection. Predictors included in our models relate to our main hypothesis (i.e., primary activity of landholders) and alternative factors explaining spatiotemporal adverse selection [i.e., 1), whether landowners are local or absentee, 2) parcel size, and 3) type of land use allowed at the site] and additive effects of these factors.

| Hypothesis | D.F | logLik | AICc | Delta AICc | Model weight |
| --- | --- | --- | --- | --- | --- |
| L.U. plan type + Local/Absentee | 15 | -927.59 | 1885.80 | 0.00 | 0.95 |
| L.U. plan type + Local/Absentee + Size | 18 | -927.33 | 1891.60 | 5.76 | 0.05 |
| L.U. plan type | 12 | -942.59 | 1909.60 | 23.79 | 0.00 |
| L.U. plan type + Size | 15 | -942.18 | 1915.00 | 29.18 | 0.00 |
| Primary activity | 21 | -958.96 | 1961.20 | 75.36 | 0.00 |
| Primary activity + Size | 24 | -958.80 | 1967.20 | 81.41 | 0.00 |
| Local/Absentee | 6 | -989.83 | 1991.80 | 105.96 | 0.00 |
| Size + Local/Absentee | 9 | -988.84 | 1995.90 | 110.09 | 0.00 |
| Null model | 3 | -1004.58 | 2015.20 | 129.38 | 0.00 |
| Size | 6 | -1003.20 | 2018.50 | 132.69 | 0.00 |

Abbreviation codes:

We ranked each model based on Akaike’s information criterion adjusted for finite sample sizes (AICc). We considered models with the lowest AICc as the most parsimonious. Modeling and data manipulation were done in program R (R version 3.4.2, R Development Core Team, 2008) with the package “nnet” for building MLRM (Venables and Ripley, 2002), and package “MuMIn” for model selection (Barton, 2018). With these models, we identified land-use plans, local versus absentee landownership, and parcel size as important predictors of adverse selection. Determining contributions of different types of landholders to adverse selection is important for targeted revision of policy and program implementation. Therefore, post-hoc we examined the relationship between primary activity of participants and 1) type of land-use plans they submitted, 2) local versus absentee landownership, and 3) parcel size. Seventy-five percent of program participants enrolled land smaller than 500 ha (median = 146 ha ± SD = 9,439.9 ha); parcel sizes in the upper quartile ranged from 500-150,000 ha (median = 1,310 ha ± SD = 18,564.5 ha). Therefore, we also tested whether the odds of adverse selection differed between small and large parcels. We built a MLRM with spatiotemporal adverse selection categories as response variables and a binary predictor variable for parcels smaller or larger than 500 ha.

## 3. Results

Primary activity of participants was not the best predictor for adverse selection (Table 1). Proposed land use under PES, whether participants were local or absentee landowners, and the size of enrolled land were stronger predictors of spatial and temporal enrollment patterns (Table 1, Fig. 2, Fig. 3), although these factors were correlated with the primary activity of participants (Table S-1). On average, agricultural suitability and contract length were lower for parcels with land-use plans for restoration and conservation than land parcels with silviculture, non-timber forest products, and silviculture plans, resulting in lower odds of adverse selection in landscapes subject to resource use than in landscapes under restoration and conservation (Fig. 2, Table S-2). For example, compared to land parcels with conservation plans, land parcels with silviculture - silvopastoral land-use plans had 193% lower odds of incurring combined temporal and spatial adverse selection (i.e., the WORST scenario of lands with low agricultural potential enrolled for short periods of time), 252% lower odds of incurring high temporal adverse selection, and 138% lower odds of experiencing high spatial adverse selection. Similarly, parcels with land-use plans for silviculture or non-timber forest products had 151% lower odds of incurring the WORST scenario, 133% lower odds of incurring high temporal adverse selection, 76% lower odds of incurring high spatial adverse selection than parcels with conservation plans (Fig. 3, Table S-2).

**Fig. 3.**
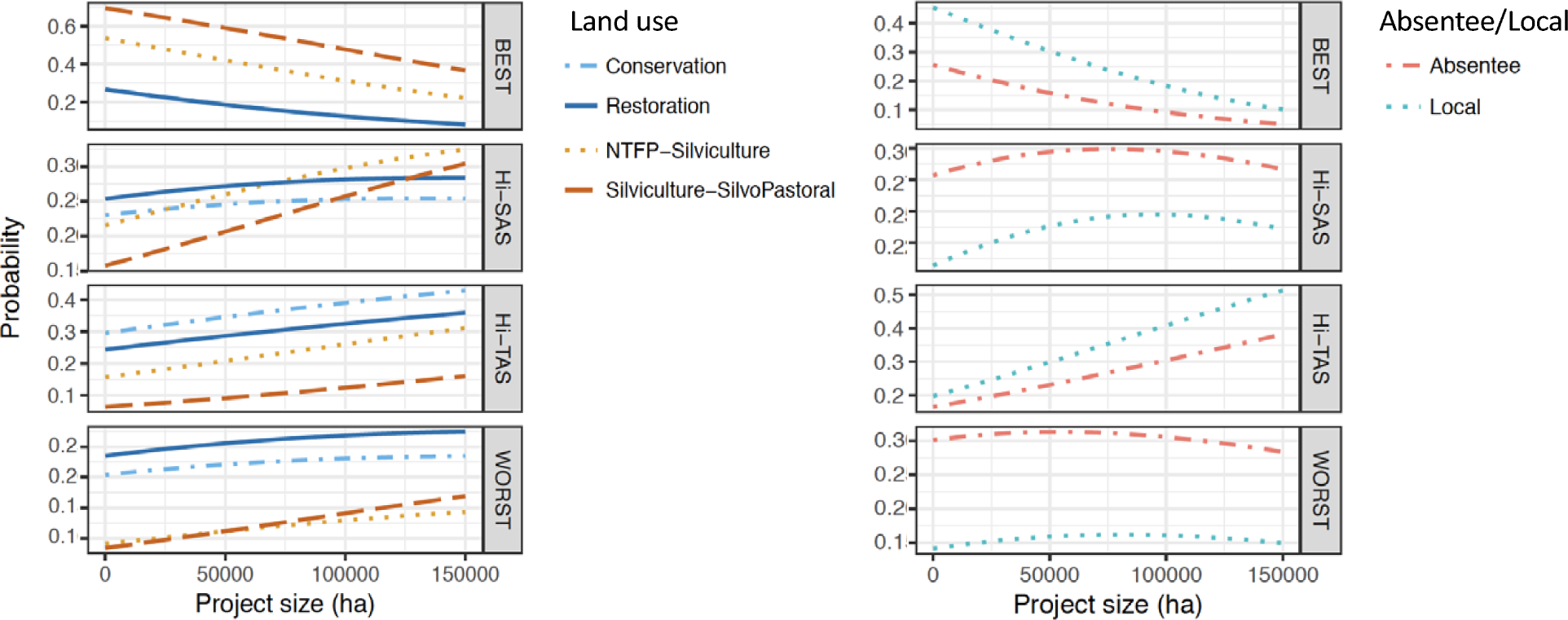
Relationships between probabilities of incurring adverse selection under WORST, Hi-TAS, Hi-SAS, and BEST and project size for (A) different land-use plans under PES and (B) local or absentee landownsers.

Governmental and non-governmental organizations (NGO) organizations and indigenous and campesino communities enrolled a larger proportion of their projects, and a greater total amount of land, with conservation and restoration plans (Tables S-3 and S-7), as compared to participants who were directly engaged with production activities or non-agricultural businesses (i.e., agriculture, ranching, silviculture, legal and real estate firms, and non-agricultural business; Tables S-3 and S-7). However, the number of conservation and restoration plans, as well as the total number of projects, submitted by participants engaged in production activities and non-agriculture businesses were much greater than the number submitted by governmental and NGO organizations and indigenous and campesino communities (Table S-3). Thus, 71% of the total conservation and restoration projects were submitted by participants that were engaged in agricultural and cattle ranching activities (44%) or non-agricultural businesses (27%, Table S-3). For all participants, lands enrolled with conservation and restoration plans had lower agricultural potential and shorter enrollment times compared to lands enrolled in production-oriented land-use plans (Tables S-4 and S-5).

Lands enrolled by absentee landowners had lower agricultural suitability than land enrolled by local land owners, and contracts were shorter for absentee landowners (Fig. 2). As a result, absentee landowners had 143% higher odds of incurring spatial and temporal adverse selection, 51% higher odds of Hi-TAS, and 96% higher odds of Hi-SAS than local landowners (Table S-2). Absentee versus local status of land owners was correlated with their primary activity (Table S-1). All indigenous and campesino communities were local landowners. The proportion of landowners dedicated to silviculture and cattle ranching was greater for local than absentee landowners (Table S-6). Landowners with a legal or a real-estate related practice were more common among absentee landowners (Table S-6). Participants dedicated to agriculture, businesses outside the agriculture industry, and governmental and NGO organizations were evenly distributed among local and absentee landowners. The probability of enrolling in BEST and Hi-TAS increased with decreasing size of parcel enrolled and the probability of enrolling in WORST and Hi-TAS increased with increasing parcel size, although these relationships were not significant (Fig. 3, Table S-2). However, parcels >500 ha had 155% higher odds of incurring both spatial and temporal adverse selection (WORST), 116% higher odds of Hi-TAS, and 204% higher odds of Hi-SAS, in comparison to land <500 ha (Fig. S-1).

## 4. Discussion

Market-based strategies like PES have the potential of becoming important policy tools to conserve ecosystem services and reduce deforestation. However, overcoming adverse selection, a key limitation to the effectiveness of PES programs, remains a critical challenge (Ferraro, 2011; Arriagada et al., 2012; Pagiola et al., 2016; Alix-Garcia et al., 2015; Börner et al., 2017; Wunder et al., 2018). We hypothesized that the primary activity of participants, particularly whether participants were engaged in agriculture, would be a strong predictor of spatial and temporal adverse selection in forested landscapes undergoing conversion to agriculture. However, other factors related to whether landowners were local or absentee, the type of land-use plan for the property under the PES program, and the size of the parcel of land enrolled were more important. Local landowners that submitted land-use plans related to production activities (non-timber forest products, silviculture, or silvopasture) were the least likely to incur spatiotemporal adverse selection (Fig. 2, Fig. 3). Absentee landowners with conservation and restoration projects were the most likely to enroll land with low agricultural potential and to enroll for short periods of time. Thus, under the current program structure, land uses that allow higher potential financial earnings and lands owned by local landowners are better suited to avoid spatial and temporal adverse selection as compared to land uses that have lower potential earnings, such as conservation projects, and are owned by absentee landowners.

Absentee landowners generally submitted land with lower agricultural potential for shorter time periods than local landowners across all categories of participant activity (Fig. 2). Other studies have found that absentee landowners both own land with lower productive potential and are less engaged in land-management activities than local landowners (Miranda et al., 2003; Arriagada et al., 2009). In Costa Rica’s PES program, absentee landowners are wealthier than local landowners and a main motivation for enrollment is lack of a better alternative use of land (Miranda et al., 2003; Arriagada et al., 2009). Studies have shown that absentee landowners of forest lands in the northeastern United States are less interested in forest management or conservation than local landowners (Petrzelka et al., 2013), characteristics that might be shared with absentee landowners in the Chaco region (le Polain de Waroux et al., 2017; this study).

In our study area, governmental and NGO organizations and indigenous and campesino communities submitted a large proportion of their projects with conservation and restoration plans (Table S-3). In Chaco, as in other regions, public lands and indigenous territories often are located in marginal lands with low agricultural productivity (Korovkin, 1997; Kareiva et al., 2007; de la Cadena, 2010; Marinaro et al., 2017; Murdock et al., 2007; but see Sims and Alix-Garcia, 2017). If most conservation and restoration plans in our study were submitted by governmental and NGO organizations and indigenous and campesino communities, a priori we might expect an association of spatial adverse selection with conservation and restoration plans. However, the total number of projects submitted by these groups was small, and landowners engaged in economic activities with high earning potential, such as agriculture or ranching, submitted most conservation and restoration projects (Table S-3). The agriculture potential of lands in conservation and restoration projects submitted by these participants was, on average, 12% lower than the agricultural potential of lands submitted as projects with collection of non-timber forest products, silviculture, or silvopasture (Table S-4).

Contract duration for conservation and restoration plans was less than half the median length of contracts with silviculture-silvopastoral land-use plans, the land use with the longest contracts time. In recent years, some participants have pushed for a re-categorization of land under high land-use restrictions (i.e., yellow and red zones) to the category that allows land conversion and land-use practices such as agriculture (i.e., green zone; le Polain de Waroux, 2017). Thus, some participants may enroll for short time-periods in conservation and restoration projects while waiting for downgrading of their land towards a zone with lower land-use restrictions. Shorter contracts also allow landholders to adapt in the face of changing market conditions by providing the flexibility to change land uses upon contract end (Roberts and Lunowski, 2007; Engel, 2016). In our study area, a short-term contract with a conservation or restoration plan also could be a strategy of some private landowners who are not currently involved in production activities but may wish to use these lands for production in the future (e.g., lands in green and yellow zones). In the case of Chaco, short enrollment contracts do not necessarily imply deforestation. The Argentine PES program allows re-enrollment of participants (García Collazo et al., 2013). Short term enrollment followed by re-enrollment could be used as a strategy to obtain inflation-adjusted payments if payments have the potential to increase over time.

Small parcels enrolled in PES in the Chaco have a lower likelihood of incurring adverse selection than large parcels. This pattern might be explained, at least partially, by disparities in opportunity costs associated with different size parcels. Land owners may be more likely to enroll small parcels for long periods of time and in areas with high agricultural suitability because of potentially lower profit margins from agriculture on small parcels. All else being equal, larger farms have a smaller cost of production per unit of land than smaller lands (e.g., returns to scale) and thus higher profit margins that must be offset by PES payments (Duffy, 2009). However, in the Chaco, enrolled parcel size may not always reflect opportunity costs because these parcels may be part of much larger landholdings that are managed as a unit. Understanding factors that limit enrollment of large parcels, particularly those with high agricultural suitability, deserves more attention. Presently, in the Chaco and elsewhere, large tracts of lands, which may be the most valuable from a biodiversity conservation perspective, likely are the most difficult to protect from deforestation through long-term enrollment in the PES programs (Salzman et al., 2018).

## 4.1 Implications for natural resource policy and conservation

The Forest Law through land-use zoning and PES aims to improve and maintain ecological and cultural processes that occur in native forest, and enrich, conserve, and restore Argentine native forests (García Collazo et al., 2013). Perhaps the most startling trend from this study for long-term biodiversity conservation is that only 10% of the parcels >500 ha were enrolled in areas under high threat of deforestation (i.e., areas with high agricultural potential), compared to 90% of the smaller parcels enrolled in these areas. If small patches of fragmented forests are disproportionately conserved with PES, long-term species persistence, and the ecological services offered by these species, will be hampered. This is especially true for wide-ranging wildlife species dependent upon large expanses of habitat and dispersal-limited species that cannot move across fragmented landscapes (Fahrig, 2003; Cushman, 2006; Quiroga et al., 2016). Small fragments also consist primarily of edge habitat where altered microclimate can induce high tree mortality and degrade the forest within the fragment, resulting in further loss of forest and forest-dependent species over time (Laurance et. al., 2011).

Enrollment of conservation and restoration plans in areas with low deforestation pressure and for short time-periods challenges the long-term effectiveness of this PES program. If government, indigenous people and local community lands have lower agricultural potential than lands dedicated to agriculture, and thus lower potential for deforestation, then spatial adverse selection is unavoidable if these lands are enrolled in PES. At current payment levels and based only on looking at adverse selection, the conservation value of investing PES funds in these lands may be questionable. However, such funds could be key in supplementing small budgets for historically marginalized communities and public land management. This is especially important in cases like in our study area where most of the total land extension was submitted by government and NGO entities and by campesinos and indigenous people (Table S-7). Furthermore, if enrollment times could be increased, commitments to PES potentially could help retain public ownership of lands during periods when government authorities support sales of public land (Schmidt, 2012).

Enrolling lands with high opportunity costs for long periods of time in PES is challenging, and few models exist for PES programs that address rapid expansion of industrial-scale agriculture and prevent deforestation. Despite this, PES-induced forest conservation has been observed in projects that aim to increase adoption of production activities that provide for carbon sequestration and some benefits for biodiversity conservation rather than adoption of stronger conservation practices with limited potential for financial gains beyond direct payments (Wunder et al., 2008; Bohlen et al., 2009; Zabel and Engel, 2010; Arriagada et al., 2012; Alix-Garcia et al., 2015, Pagiola et al., 2016, Jayachandran et al., 2017). These two scenarios mirror land-use categories under the Forest Law that allow for lower-impact economic activities like silviculture or silvopasture, (e.g., yellow zone) and land-use categories that prohibit these practices but encourage conservation and restoration projects (e.g., red zone) of the Argentine PES. In a PES program in Mexico, the greatest additionality was found when land close to park edges enrolled in PES (Sims and Alix-Garcia, 2017). Similarly, the greatest additionality in the Argentine PES may occur where enrolled lands are in yellow zones adjacent to protected areas or other lands in the red zone.

In the short term, PES in the Chaco may be most successful in enrolling private lands with projects focused on land uses with earning potential (e.g., silviculture or silvopasture) in areas with intermediate protection levels (i.e., yellow zones). These land uses reduce opportunity costs and increase the chances that a given payment will match or exceed minimum acceptable payment threshold for supplying environmental services (Börner et al., 2017). In our study area, a larger proportion of land in the red zone is public land (in comparison to other land-use zones), occurs on lands with low agricultural suitability, and already has some form of protection. In contrast, most lands in the yellow category are private landholdings with higher agricultural suitability. Because regional land-use plans prohibit deforestation but allow limited production activities in the yellow category, payments constitute incremental income above income generated by low-intensity production activities (e.g., cattle ranching) rather than a replacement for all income generated by conversion to large scale agriculture.

In the long term, PES projects with conservation and restoration plans, which restrict production activities on private lands, should provide the best management strategies to accomplish the goals of the Argentine PES. Private lands have high potential for land conversion or degradation with production activities, and these land-use plans offer considerably more protection than plans that allow production. For example, a silvopasture approach called “Forest Management with Integrated Cattle Ranching” (MBGI-Manejo de Bosques con Ganadería Integrada), which allows for tree removal and planting of introduced grasses, is being promoted as a compatible strategy for both biodiversity conservation and cattle ranching (FVS, 2016). MBGI is widely supported across some government and non-government organizations, as well as by private land-owners, and could be used in the largest stretches of remnant Chaco forest (i.e., yellow and green zones). However, whether MBGI will meet the Forest Law’s objectives of conserving biodiversity is unknown. Specialists have raised concerns about potential negative effects on biodiversity such as increases in forest fragmentation, reduction in the quantity and quality of wildlife habitat, and synergistic increases in hunting pressure (FVS, 2016). These specialists also point out that implementing corridors and promoting sustainable hunting practices could mitigate some effects of silvopasture activities such as MBGI (FVS, 2016). However, the success of such a mitigation strategy still will depend on having extensive forest blocks to link with corridors, suitable habitat within corridors, and new strategies for managing hunting. Increasing incentives for long term enrollment of threatened land under conservation and restoration projects might be a more parsimonious strategy to meet the program’s goals.

The degree to which economic development and conservation goals can be met simultaneously with PES will depend on the compatibility of silvopasture practices, as well as extraction of timber and non-timber forest products, with forest and biodiversity conservation. Considerable effort has focused on development of management options for sustainable use of humid and seasonally dry tropical forests that meet conservation goals (Putz et al., 2008). However, much less is known about compatible management for semiarid, subtropical forests such as Chaco (Trigo et al., 2017). The challenges are significant, ranging from climate extremes that limit forest productivity and necessitate long rotation cycles for ecologically sustainable harvest to severe ecosystem degradation from decades of unmanaged grazing (Grau et al., 2008). Furthermore, although the largest amount of land in the Chaco is in the yellow category slated for sustainable use, some very critical pieces for long-term regional conservation, such as forest corridors that link protected areas and safeguard riparian areas, occur on private lands in the red category where use is restricted (Núñez-Regueiro pers. obs.). If PES cannot engage these landowners in long term contracts that conserve these regions, then the limits of PES need to be recognized and other strategies need to be identified and implemented.

## 5. Conclusions

Maintaining native forests with low human intervention is critical for supporting the world’s ecosystem services, however wild places continue to disappear (Potapov et al., 2017). Developing strategies to encourage conservation and restoration projects that avoid adverse selection under PES in forested areas is fundamental to the success of these programs in addressing forest loss. Understanding predictors for spatial and temporal enrollment patterns may help improve effectiveness of PES programs, incentivize protection of forests, and address critical environmental challenges such as deforestation and climate change (Ferraro, 2011; Alix-Garcia et al., 2015; Chazdon, 2017).

Under the current PES structure in Chaco, the most feasible means of achieving enrollment for threaten lands is through land-use activities that simultaneously promote sustainable forest use and conservation (e.g., silvopasture), as opposed to practices that restrict land use to conservation or restoration alone, and by encouraging active participation of local landowners. However, ecological impacts of land use on biodiversity, as well as forest cover, need to be assessed to recommend land-use practices under PES. Furthermore, although PES contracts for production-related land-use plans were less likely to be adversely selected than conservation-oriented plans, enrollment periods for all participants and land uses were short compared to the timeframe needed for sustainable forest management and biodiversity conservation. Providing bonus lump sum payments for long-term contracts or linking payments to commodity prices could incentivize long-term contracts by counteracting decreasing marginal benefit for landowners remaining in the program for long time-periods (Juutinen et al., 2014). Additionally, offering the highest-tiered payment levels for conservation and restoration lands in yellow and green zones could maximize the program’s additionality by incentivizing land-use practices highly aligned with the program’s goals in areas with high threat of land-use conversion.

Our results also indicate that large land parcels are least likely to achieve long-term enrollment in highly productive areas, which could accentuate already alarming levels of habitat fragmentation and biodiversity loss (Núñez-Regueiro et al., 2015; Quiroga et al., 2016). Thus, strategies need to be developed and implemented that promote enrollment of large parcels of forest under high threat. One potential approach is offering payments proportional to the land’s agricultural value. Spatial targeting of PES also is fundamental to avoid spatial adverse selection and to increase the overall size of contiguous protected forests through enrollment of adjacent land parcels and parcels bordering protected areas. Finally, monitoring PES program’s performance under an adaptive management framework, as well as identifying where conservation objectives and the PES program are poorly matched from the outset, will be key for land management that supports the goals of PES and for improving conservation outcomes of PES (Sims et al., 2014; Alix Garcia et al., 2015).

